# Molecular insight into Nickel(II) binding by Her2-specific IgE: a possible mechanistic insight into the pathogenesis of Type I nickel hypersensitivity

**DOI:** 10.1101/2020.07.14.203539

**Authors:** Chinh Tran-To Su, Wai-Heng Lua, Jun-Jie Poh, Wei-Li Ling, Joshua Yi Yeo, Samuel Ken-En Gan

**Affiliations:** Experimental Drug Development Centre, Agency for Science, Technology and Research (A*STAR), Singapore 138670; Antibody & Product Development Lab, Bioinformatics Institute, A*STAR, Singapore 138672; p53 Laboratory, A*STAR, Singapore 138648

**Keywords:** nickel allergy, IgE, antibody VH-VL families, Pertuzumab, Trastuzumab, glutamine, histidine, Type I hypersensitivity

## Abstract

Nickel (Ni) allergy has been reported in contact dermatitis Type IV (Ni-specific T cells mediated) and asthmatic Type I (IgE-mediated) hypersensitivities. Associations between the two hypersensitivities have been found in some patients, but the underlying mechanisms remain enigmatic. Using Her2-specific IgEs as models, we found additional binding to Ni-NTA without observable changes in binding to Her2 and that glutamine, together with the canonical Ni^2+^-binding histidine, could form Ni^2+^ binding signatures. This mechanism may underlie Type I hypersensitivity in the selection of anti-Ni^2+^ IgEs. This mechanism may also underlie Type IV hypersensitivity and the interaction of immunoglobulin proteins with other heavy metal ions. Our findings shed light to how Ni hypersensitivities can occur and how they can be avoided in therapeutics design, or even incorporated for biotechnological purification purposes.

## INTRODUCTION

Nickel allergy is one of the most common metal allergies (Ahlström et al., 2019) given the inevitable daily exposure in common metal alloys present in household products and even in food (Flyvholm et al., 1984). Nickel allergy is usually reported as contact dermatitis of Type IV hypersensitivity and to a lower incidence, Type I hypersensitivity. While Type IV hypersensitivity is mediated by hapten-specific T cells (Aparicio-Soto et al., 2020; Büdinger and Hertl, 2000; Büdinger et al., 2001; Saito et al., 2016; Schmidt et al., 2010), Type I nickel hypersensitivity is IgE-mediated (Bennike and Foss-Skiftesvik, 2018; Dolovich et al., 1984; Kolberg et al., 2020; Malo et al., 1982; Nieboer et al., 1984; Novey et al., 1983; Saluja et al., 2016; Sastre et al., 2001; Shirakawa et al., 1990). Increasingly, associations between Type IV and Type I hypersensitivities to nickel in populations are reported (Gelardi et al., 2017; Kolberg et al., 2020), with evidence of both type I and IV hypersensitivity occurring within the same patient (Walsh et al., 2010). It remains unclear if the two allergy types share similar underlying mechanisms.

Structurally, similar topologies exist between the variable regions (both framework regions or FWRs and complementarity-determining regions or CDRs) of antibodies such as IgE and the T cell receptor (TCR). Elucidating the mechanisms of Ni^2+^ binding in type I hypersensitivity may provide clues also to type IV hypersensitivity that involves interactions between the human leukocyte antigen (HLA)/peptide complex and TCR.

Occupational asthma is the most commonly reported type I nickel hypersensitivity, and results from prolonged exposures to nickel (Brera and Nicolini, 2005; Bright et al., 1997; Dolovich et al., 1984; Malo et al., 1982; Malo et al., 1985; Nieboer et al., 1984; Novey et al., 1983; Sastre et al., 2001; Shirakawa et al., 1990) leading to Ni specific IgEs.

The Ni^2+^ ion binds to histidine residues and this interaction is commonly exploited in protein purification methods involving the fusion with histidine tags (Hemdan et al., 1989; Knecht et al., 2009; Lebrette et al., 2014; Porath et al., 1975; Sudan et al., 2015), underlies the interaction of the major histocompatibility complex (MHC) with certain peptides (Driller et al., 2019; Gamerdinger et al., 2003; Vollmer et al., 1999), as well as in several proteins such as Ηpn-like protein in *H. pylori* and other bacteria (Zeng et al., 2011; Zeng et al., 2008). The same interaction is found in Fcγ (Hale and Beidler, 1994; Rosen et al., 2014) and in TCRs of Type IV contact dermatitis patients, where increased Ni^2+^-binding histidine residues were found (Aparicio-Soto et al., 2020). A single domain VHH antibody engineered to have three added histidine residues binding Ni^2+^ in the hypervariable loops also showed disruption of antigen binding (Fanning et al., 2014).

While less frequent than in type IV, nickel could elicit type I hypersensitivity through “non-antigenic” mechanism via IgE (Malo et al., 1985) and was postulated to act as a mast cell discharger (Walsh et al., 2010). It is possible that Ni^2+^ binding may involve mechanisms apart from CDRs alone, requiring a whole antibody holistic investigation (Phua et al., 2019) given that the various regions of the antibody interact together to affect antigen binding (Lua et al., 2018; Su et al., 2018) and even receptor engagement (Ling et al., 2018; Lua et al., 2019), with the possibility of these regions coming synergistically to form histidine rich areas to bind Ni^2+^.

In our previous work (Lua et al., 2019), we found our recombinant IgEs of varying VH regions to bind FcεRIα differentially. Further analysis found that some of the IgEs also bound to Ni-NTA without compromising the binding of the antigen Her2. In this present work, we investigate the possible mechanisms of this Ni-IgE interaction that has relevance in purification purposes, in avoiding unwanted side effects for therapeutic antibodies, and in nickel type I and type IV hypersensitivities.

## RESULTS

### Effect of Antibody VHs on nickel binding

Recombinant Pertuzumab and Trastuzumab IgE and IgG1 variants with VH families 1–7 paired with their respective wild-type Pertuzumab or Trastuzumab Vκ1 were analyzed with respect to their interactions with the nickel-attached nitrilotriacetic acid biosensor (Ni-NTA) using bio-layer interferometry on the Octet RED96 system. Experiments of direct NTA (without Ni^2+^) with the Pertuzumab VH5 IgE ruled out interactions with the NTA alone (Supplementary Figure S1).

All the Pertuzumab IgE VH family variants (except for the VH3 and VH6) had measurable equilibrium dissociation constants (KD) with the Ni-NTA sensor. The KD of the Pertuzumab VH5 IgE variant was the smallest (strongest interaction, Figure 1A). On the other hand, all the Pertuzumab IgG1 variants, except for Pertuzumab VH5 IgG1, had poor responses in the measurements (Figure 1B). Thus, the change of the C-region (CHs) from IgE to IgG1 isotypes in all the Pertuzumab variants, including in VH5, diminished interactions to the Ni-NTA. On the other hand, for Trastuzumab VH3 and VH5, variants, switching from IgE to IgG1 improved the KD of Ni-NTA binding, demonstrating Ni-binding to be contributed by the various regions of the antibodies.

**Figure 1:**
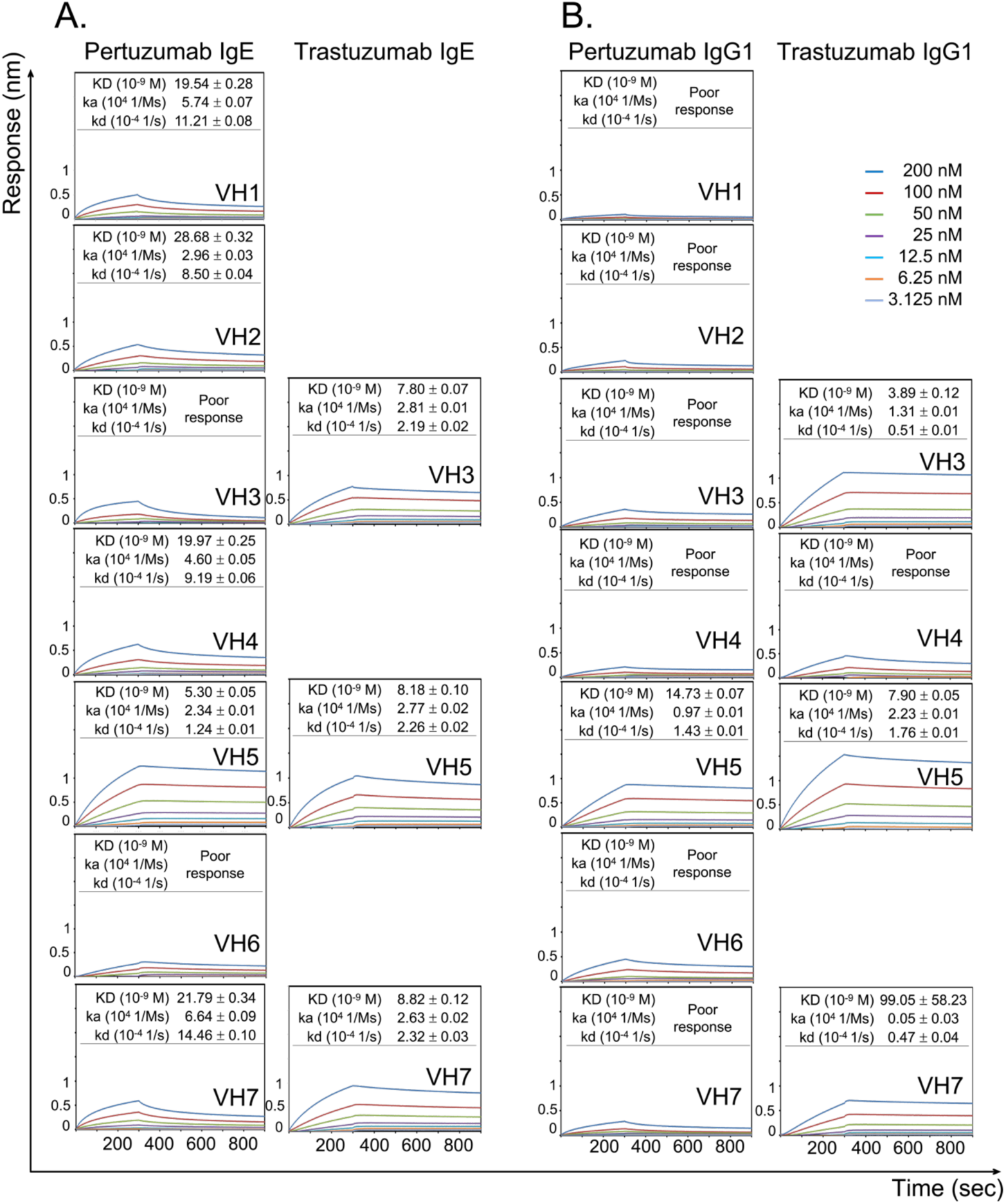
Measurements of dissociation equilibrium constants (KD) of Pertuzumab and Trastuzumab IgE and IgG1 of different VH families paired with the respective Pertuzumab and Trastuzumab Vκ1 light chain to show their interactions with the Ni-NTA biosensor. The measurements were performed at various concentrations from 200 nM to 3.125 nM of the antibodies. Values of KD (M), ka (1/Ms), and kd (1/s) were measured and calculated using the Octet RED96 system. The X-axis depicts the time (in seconds) while the Y-axis depicts the binding responses (nm). All the experiments were conducted in at least duplicates. “Poor response” is determined for the cases where the responses are less than 0.1 nm for at least three concentrations. Those variants that could not be successfully produced in our recombinant transient transfections are left blank.

Within the respective Pertuzumab and Trastuzumab variants, the various VH1–VH7 variants, while paired to the respective Pertuzumab or Trastuzumab Vκ1 light chain counterparts, showed more pronounced Ni-NTA KD measurements (Figure 1). These variants bear the same respective CDRs within each model. Among the Pertuzumab variants, the Pertuzumab VH5 IgE and IgG1 variants showed the lowest (strongest) KD ∼5.30 nM and ∼14.73 nM, respectively. Changing the CDRs to Trastuzumab in the VH family variants, gave lower KDs (better binding) for VH3 and VH7 IgE and also IgG1 (including VH5 IgG1), although the reverse was observed for Pertuzumab VH5 IgE. As described previously, only limited Trastuzumab VH variants could be produced in our expression system (Lua et al., 2019) due to various reasons (unpublished).

### Glutamine, in addition to Histidine, binds Ni-NTA

The histidine (H) and glutamine (Q) counts were found to be significantly varied in the V-region FWRs and CDRs of the Pertuzumab and Trastuzumab variants (Supplementary Table S1). Although histidine is a known binder of Ni^2+^ (Hale and Beidler, 1994; Hemdan et al., 1989; Rosen et al., 2014), glutamine was also reported to have long-range effects on Ni^2+^ binding in several proteins such as Hpn-like proteins (Chiera et al., 2013) or directly bind Ni^2+^ e.g. in HLA-DR52C (Wang and Dai, 2013) and in a single domain antibody (Fanning et al., 2014).

Trastuzumab VH3, VH5, and VH7 IgE had a glutamine stretch that was found in docking simulations to be predominantly involved in the Ni-NTA binding (Figure 2 and Supplementary Figure S2), as opposed to 0.4% (Q) vs 59.23% (H) of total 8,054 Ni^2+^ interactions retrieved from 1,810 protein-Ni^2+^ complexes found in Protein Data Bank (as of June 2020). This suggests that residue content alone (of histidine or glutamine) do not predict nickel binding, but the structural arrangement of these residues.

**Figure 2:**
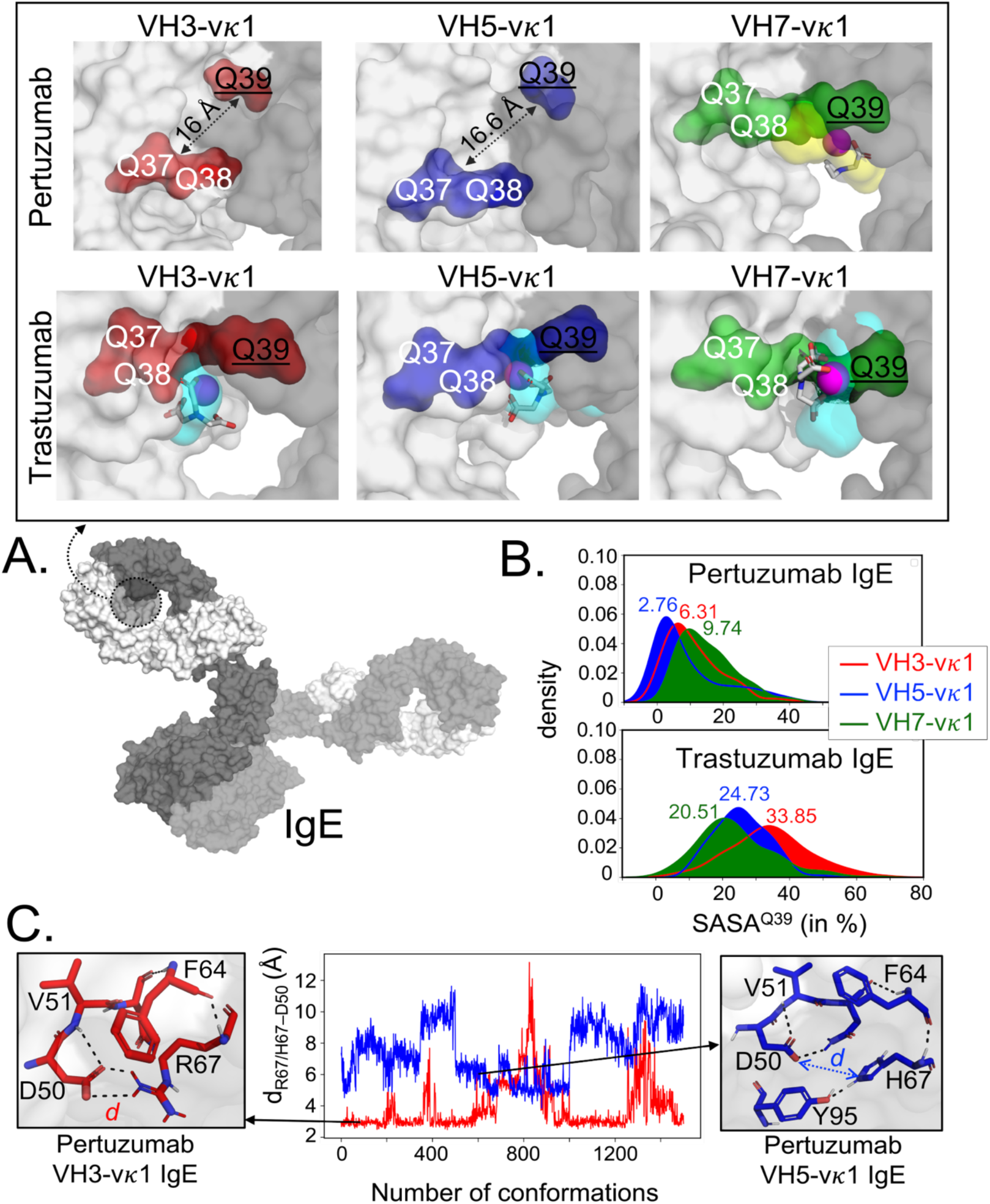
Structural analyses of Pertuzumab and Trastuzumab VH3, VH5 and VH7 IgE variants to show glutamine, aside from histidine, to bind to Ni-NTA. **(A)**. Model of full length IgE (light chain in white surface and heavy chain in gray surface) reveals the formed pocket at the VHx–Vκ1 interface (dashed circle). The Ni-NTA bound region of the variants VH3 (red), VH5 (blue), and VH7 (green) shows the glutamine (Q) stretch constituted by the VH Q39 (underlined) and Vκ1 Q37-Q38 to form a stable cryptic pocket (capacity of which is shown in cyan transparent surface) in the Trastuzumab IgE variants or a transient non-cryptic pocket (yellow transparent surface) in the Pertuzumab VH7. The bound Ni-NTA is shown in magenta spheres and interacts at the pocket in both Pertuzumab and Trastuzumab. **(B)**. Relative solvent accessible surface area (SASA) of the Pertuzumab and Trastuzumab VH Q39 residue in all the variants to demonstrate the deeply buried Q39 (SASA< 10%) of the Pertuzumab variants. **(C)** Distance *d* between residue 67 (R67 in VH3 or H67 in VH5) and heavy chain CDR2 D50 of Pertuzumab VH3-Vκ1 IgE (red) and VH5-Vκ1 IgE (blue) during the simulation. Structural visualizations were generated using PyMOL 2.3.2 (Schrodinger, 2019).

Structures of full-length Pertuzumab and Trastuzumab IgE of VH3, VH5, and VH7 families paired with their respective Vκ1 light chains were modelled and the different conformations of their Fab (containing VHx-Vκ1, Cκ, and Cε1) regions were used in blind dockings with a parameterized Ni-NTA molecule to explore the regions or cavities for the interactions. The Ni-NTA was found to predominantly bind to residues Q38 (Vκ1 FWR2) and Q39 (VH3, VH5, and VH7 FWR2) of the Trastuzumab IgE variants (Supplementary Figure S2). Together with Q37 of Vκ1 FWR2, the residue Q38 (Vκ1 FWR2) and Q39 (of VH3, VH5, or VH7 FWR2) form a structural glutamine stretch that shapes a stable (in the course of 3×100 ns) internal pocket right at the interface of the heavy and light chains (cyan in Figure 2A). Interestingly, these pockets were found to be cryptic (see methods and Supplementary Table S2), allowing possible open channels for the accessibility of the Ni-NTA. The cryptic pockets were only found in Trastuzumab and not Pertuzumab V-regions, pointing to CDR effects.

The Q39 of VH7 FWR2 in the Pertuzumab IgE together with Q37 and Q38 of the Vκ1 form a non-cryptic and less dense cavity (yellow in Figure 2A). A less populated Ni-NTA binding cluster was located at the Q39 (VH7 FWR2) than that of the corresponding locations in the Trastuzumab VH7-Vκ1 IgE (Supplementary Figure S2). This finding lends support from our experimental results where Trastuzumab VH7 IgE bound the Ni-NTA better (with a lower KD) than Pertuzumab VH7 IgE (Figure 1A).

There were no such glutamine-forming pockets in the Pertuzumab VH3-Vκ1 and VH5-Vκ1 IgE variants since the VH Q39 residues were deeply buried and distant from the respective Vκ1 Q37 and Q38 (Figure 2A and 2B) due to varying heavy and light chain structural arrangement. Instead, binding of the Ni-NTA at the Vκ1 Q89, VH5 H67 (in FWR3), and heavy chain CDR2 D50 of both Pertuzumab VH3 and VH5 IgEs was observed (Supplementary Figure S2). The VH5 FWR3 H67 is present in both Trastuzumab and Pertuzumab models.

We performed single point mutagenesis of H67Q (present in VH5-10 and other VH5 germlines) for both Pertuzumab and Trastuzumab VH5–Vκ1 IgE variants and found the Ni-NTA binding affinity to diminish more drastically in the Pertuzumab VH5^H67Q^ IgE than in the Trastuzumab VH5^H67Q^ IgE, with a ∼5-fold vs ∼1.5-fold increase in KD, respectively (Figure 3). Although residue H67 is present in both antibody VH5 (FWR3), only Pertuzumab VH5 H67 was structurally involved with Vκ1 Q89 in the Ni-NTA binding, whereas the glutamine stretch Q39–Q37/Q38 of Trastuzumab VH5 IgE played the more predominant role to bind the Ni-NTA (Figure 3). The involvement of the Vκ1 Q89 in Pertuzumab VH5 IgE in another glutamine stretch is further discussed below.

**Figure 3:**
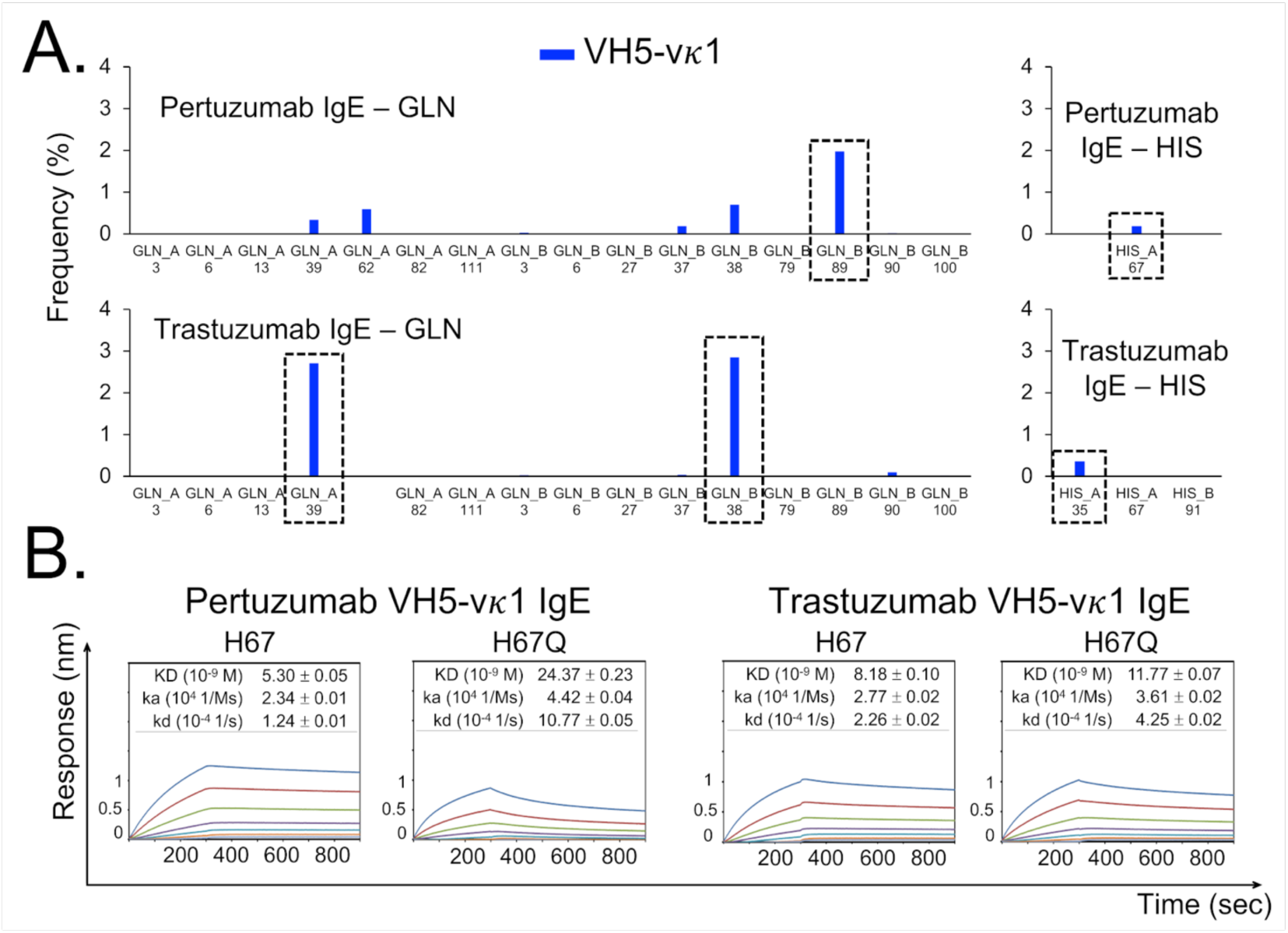
Main Ni-NTA binding region of the Pertuzumab VH5–Vκ1 IgE is at H67 of VH5 FWR3 whereas in the Trastuzumab VH5–Vκ1 IgE at the glutamine stretch containing VH5 Q39 at FWR2. **(A)** Distribution of the Ni-NTA docked conformers found at all the glutamine (GLN) and histidine (HIS) present in the V-region of the Pertuzumab and Trastuzumab VH5– Vκ1 IgE variants. The predominant binding residue locations are highlighted in dashed boxes. **(B)** Changes in dissociation equilibrium constants of the Pertuzumab and Trastuzumab VH5– Vκ1 IgE variants with mutation H67Q to the Ni-NTA sensors.

The Pertuzumab VH3 IgE had a poor response in our Ni-NTA binding measurements. (Figure 1). Structurally, the Ni-NTA docked conformers were found scattered in a few spatial regions on the Pertuzumab VH3 IgE, and mostly populated at residue D50 (located in the heavy chain CDR2) as shown in Supplementary Figure S2. Interestingly, this D50 was a part of a polar contact network that extended to the residue at position 67 of the heavy chain FWR3, i.e. R67 and H67 in Pertuzumab VH3 and VH5 IgE, respectively (Figure 2C). The D50-R67 interaction was mostly stable, possibly only allowing D50 occasional engagement of Ni-NTA. On the other hand, the D50 and H67 in the VH5 Pertuzumab IgE variant were not involved in this interaction and thus was more accessible to the Ni-NTA.

### Effect of Vκ on nickel binding

To investigate the possible effects of different Vκ families on the nickel binding, we used the Pertuzumab and Trastuzumab IgG1 variants as the main antibody isotype for light chain (Vκ) switching (see methods). Vκ1 to Vκ6 variants of Pertuzumab and Trastuzumab were individually paired with the VH3 and VH5 of both Pertuzumab and Trastuzumab IgG1. These two VHs exhibited the lowest KD in Ni-NTA binding (Figure 1).

All the Pertuzumab VH3 IgG1 variants paired with the different Vκ (except for Vκ5 that was not successfully produced) showed poor Ni-NTA binding responses (Figure 4A). Similarly, in the Trastuzumab VH3 IgG1 variants, when Vκ1 was replaced with Vκ2, Vκ3, Vκ4, or Vκ6, the Ni-NTA bindings were not measurable, and a high KD was found with Vκ5 having an increased KD of ∼12.83 nM from 3.89 nM.

**Figure 4:**
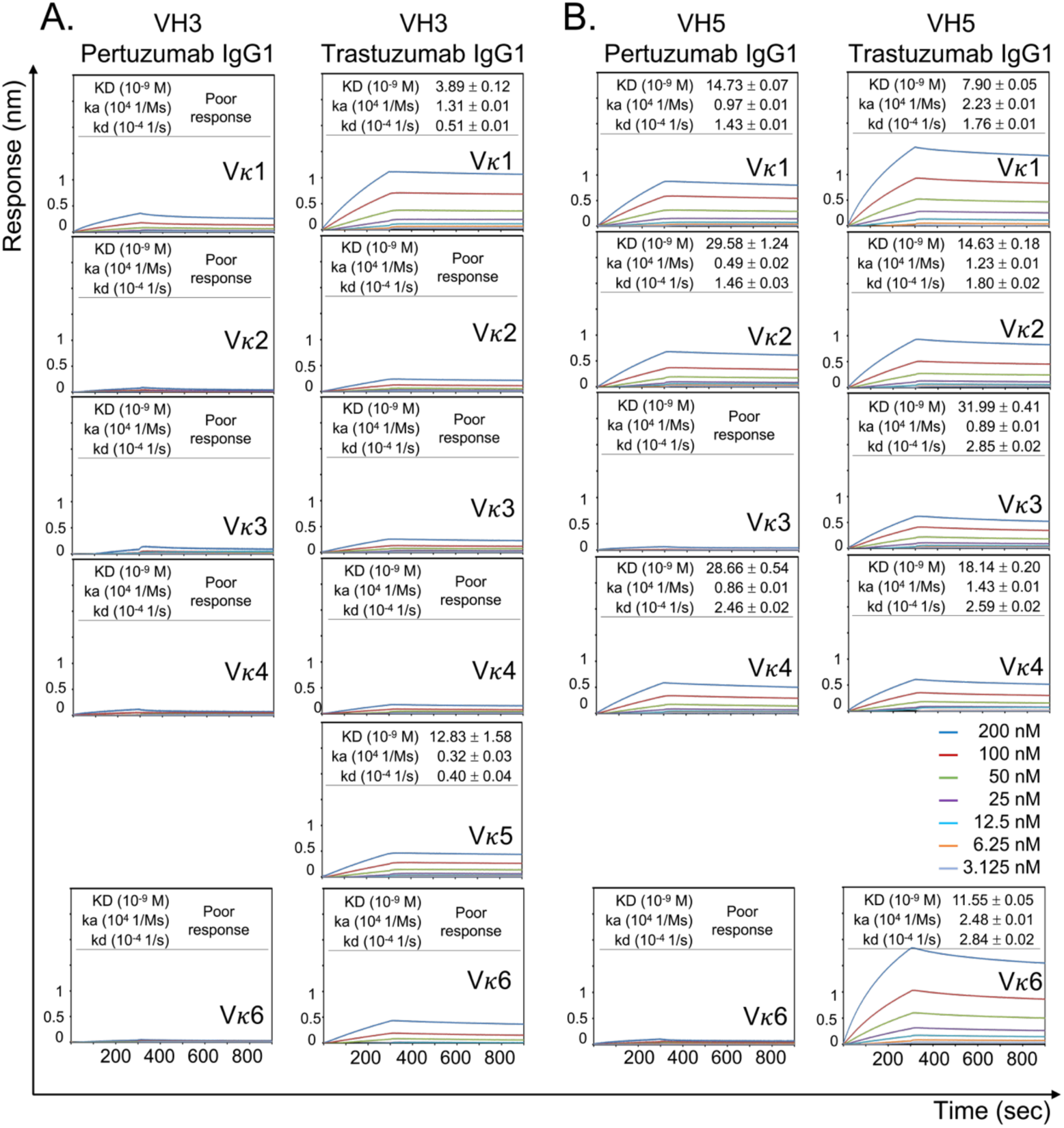
Ni-NTA dissociation equilibrium constants (KD) of Pertuzumab and Trastuzumab VH3/VH5 IgG1 variants paired with different Vκs. The experiments were performed at various concentrations from 200 nM to 3.125 nM of the antibodies. Values of KD (M), ka (1/Ms), and kd (1/s) were measured and calculated using the Octet RED96 system. The X-axis depicts the time (in seconds) while the Y-axis depicts the binding responses (nm). All the experiments were conducted in at least duplicates. “Poor response” is determined for the cases where the responses are less than 0.1 nm for at least three concentrations. Those variants that could not be successfully produced in our recombinant transient transfections are left blank.

Both the Pertuzumab and Trastuzumab VH5 IgG1 variants showed better binding to Ni-NTA than their VH3 counterparts (Figure 4B), especially when the Trastuzumab VH5 IgG1 variants was paired with Vκ2, Vκ3, Vκ4, and Vκ6 FWRs. Among the VH5 variants, except for those paired with Vκ5 that were not produced, the Pertuzumab VH5 IgG1 that paired with Vκ3 and Vκ6 had poor responses to the Ni-NTA binding measurements (Figure 4B).

Quantitatively, sequences comparisons between the Pertuzumab and Trastuzumab VH5 IgG1 (pairing respectively with the same Vκ family) showed differences only in the heavy/light chain CDRs. Therefore, results of improved Ni-NTA binding responses in the Trastuzumab VH5 IgG1 cases (Figure 4B) affirmed the CDR effects of both chains in Ni-NTA interactions. Comparing the variants with the same CDRs but varying light chain FWRs, e.g. Pertuzumab VH5–Vκ1 and VH5–Vκ3 IgG1, the opposite responses to the Ni-NTA binding were found, showing the contribution of both the FWRs and CDRs of both chains.

Within these Pertuzumab and Trastuzumab VH5 IgG1 variants, the residue contents of glutamine and histidine remained consistent between the various Vκ switches (Supplementary Table S1), showing that the residue counts alone do not determine Ni binding, but the structural arrangement between the two chains involving both CDRs and FWRs of both chains, and the CHs.

To investigate further the underlying mechanism, we modelled the full length Pertuzumab and Trastuzumab VH5–Vκ1, VH5–Vκ3, VH5–Vκ6 IgG1 and simulated the dynamics of their Fab regions.

Distance-based maps were constructed for the Pertuzumab and Trastuzumab IgG1^Fab^ of VH5-Vκ1, VH5-Vκ3, and VH5-Vκ6 variants to represent distance networks between the VH5, Cγ1, Vκ, and Cκ regions. Superimposition of these maps showed the distance differences between the regions, eliciting the conformational changes possibly caused by switching various Vκ (with the same VH5, Cγ1, and Cκ). It was shown that replacing Vκ1 with Vκ3 or Vκ6 resulted in more significant structural disruption in the Pertuzumab VH5 IgG1 variants (red color in Figure 5A), particularly in the VH5 geometry with respect to the Cκ accommodation in the Fab region. In fact, when switching to Vκ6 from Vκ1, we additionally observed shifts of distances between the Vκ6 and other domains such as Cκ and Cγ1 in the Pertuzumab IgG1 variants. On the other hand, mild changes (blue and green colors in Figure 5A to represent small differences) occurred in the Trastuzumab VH5-Vκ3 and VH5-Vκ6 IgG1.

**Figure 5:**
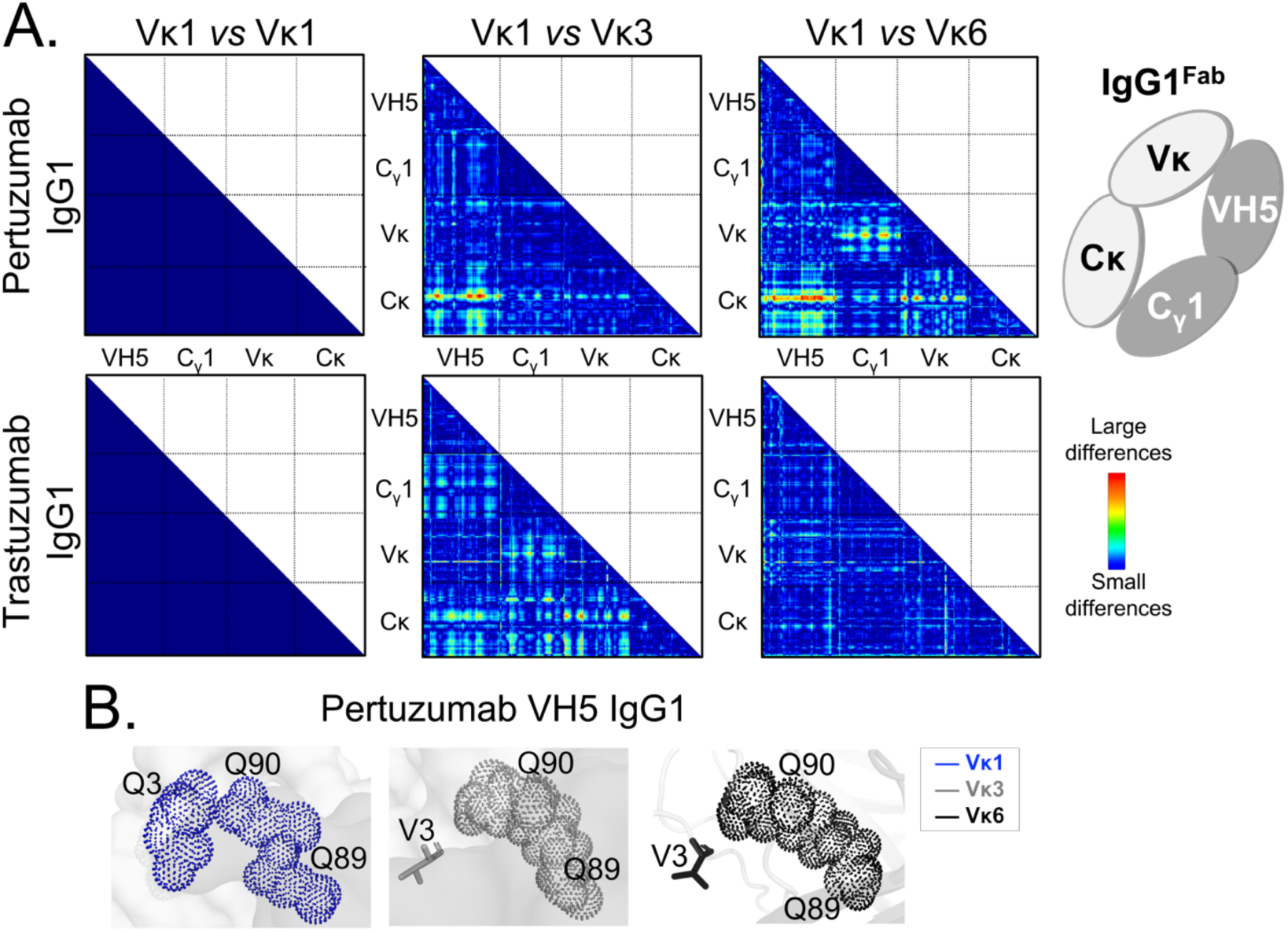
Effects of the different VH5-Vκ pairings in the Pertuzumab and Trastuzumab IgG1 variants. **(A)** Distant-based difference maps (generated using CMView (Vehlow et al., 2011)) between Fab domains of Pertuzumab and Trastuzumab VH5-Vκ3 and VH5-Vκ6 IgG1 variants (superimposed against the respective VH5-Vκ1 IgG1 as the reference) indicates more disruptive effects by the Vκ3 or Vκ6 on the Pertuzumab VH5 IgG1 variants, leading to varying changes in the heavy and light chain pairing. **(B)** Another Ni-NTA binding region of the Pertuzumab VH5-Vκ1 IgG1 variants at the formed glutamine stretch of Vκ1 Q3, Q89, and Q90. These glutamine stretches were incomplete due to the Vκ mutation Q3V in the Pertuzumab VH5-Vκ3 and VH5-Vκ6 IgG1 variants.

Results of blind dockings of the Ni-NTA to these Fab regions showed that the Trastuzumab VH5-Vκ1, VH5-Vκ3, and VH5-Vκ6 IgG1 variants predominantly bound the Ni-NTA using the glutamine stretch of VH5 Q39 and Vκ Q37-Q38 at the heavy-light chain interface, as also in Trastuzumab VH5-Vκ1 IgE (Supplementary Figure S2 and S3). Given that the heavy and light chain geometries of those Trastuzumab VH5 IgG1^Fab^ were insignificantly perturbed when switching the Vκ, the Ni-NTA binding therefore maintained. The results were also observed experimentally (Figure 4).

In line with the Ni-NTA docking results of the Pertuzumab VH5-Vκ1 IgE variant, the Ni-NTA was also found to computationally bind to the Vκ1 Q89 in the Pertuzumab VH5-Vκ1 IgG1. The frequency of these Ni-NTA interactions at the Vκ Q89 diminished in the Pertuzumab VH5-Vκ3 IgG1 and abolished in the Pertuzumab VH5-Vκ6 IgG1 (Supplementary Figure S3). Structurally, the Vκ1 Q89 is a part of another glutamine stretch together with the adjacent Vκ1 Q90 (in CDR3) and the distant Vκ1 Q3 (in FWR1). However, this glutamine cluster isincomplete in the other two Pertuzumab IgG1 variants of VH5-Vκ3 and VH5-Vκ6 due to the mutation Q3V at both the Vκ3 FWR1 and Vκ6 FWR1 (Figure 5B), likely leading to the loss of Ni-NTA binding (Figure 4B). Similar to the Trastuzumab VH5-Vκ1 IgE, the residue H67 was not structurally involved in the Ni-NTA binding in all the three Trastuzumab VH5-Vκs IgG1 variants.

### Pertuzumab VH5-Vκ1 IgE could bind multiple metal ions, possibly using both glutamine and histidine to bind Ni^2+^

To investigate further the ability of binding other metal ions beyond the Ni^2+^, we measured the interaction between Pertuzumab VH5-Vκ1 IgE variant to the NTA sensors recharged with various metal ions Cu^2+^ and Co^2+^. The Pertuzumab VH5 IgE was found to bind the metals with varying magnitudes in the order of Cu^2+^ > Ni^2+^ > Co^2+^ (Figure 6A), a trend following the Irving-Williams’ order of metal ions (Irving and Williams, 1953).

**Figure 6:**
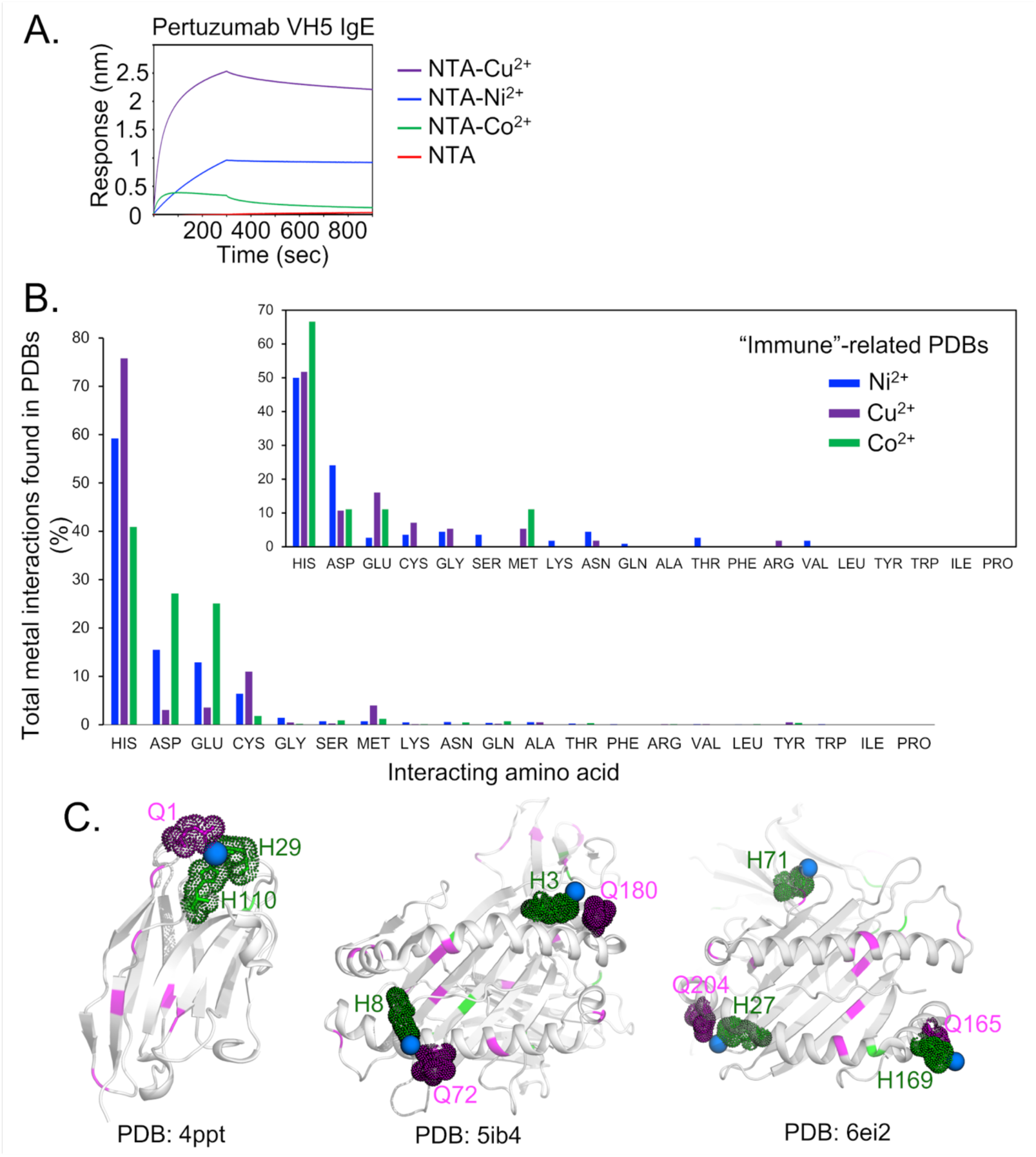
Analysis of multiple metal binding ability. **(A)** Binding responses of the Pertuzumab VH5 IgE variant to different metal ions (Ni^2+^, Cu^2+^, and Co^2+^) that were recharged on the NTA sensors. **(B)** Numbers of the metal interactions in the retrieved protein-metal complexes from Protein Data Bank. The smaller offset plot shows the metal interaction results of a subset of immune-related protein complexes found by using independently keywords: “immune”, “antibody”, “MHC”, and “TCR”. **(C)** Structural presentation of a VHH antibody complexed with Ni^2+^ (PDB: 4ppt), HLA-B*27:05 (PDB: 5ib4), and HLA-A68 (PDB: 6ei2) complexed with a peptide and Ni^2+^, where glutamine (magenta) are found to be involved in (e.g. in 4ppt) or closed to the Ni^2+^ binding region with histidine (green). The Ni^2+^ ions are represented by blue spheres.

Furthermore, our analysis of a limited number of protein-metal complexes (1409 for Cu^2+^, 1810 for Ni^2+^, and 716 for Co^2+^ retrieved from the Protein Data Bank (PDB), accessed in June 2020) showed that histidine is the most common residue that binds the multiple metals, followed by aspartate, glutamate, cysteine, methionine, serine, and glutamine (Figure 6B). However, only a small number of structures (12, 26, and 14 complexes involving the metal ions, respectively) are related to the immune system (see methods, Figure 6B, small offset). It showed that histidine remained the main binding factor to all the three metal ions whereas glutamine was found to be involved only in the binding to Ni^2+^. Notably, three protein-Ni^2+^ complexes (a single VHH antibody domain, PDB: 4ppt (Fanning et al., 2014), and two HLA-peptide complexes, PDB: 5ib4 (Driller et al., 2019) and 6ei2 (Guillaume et al., 2018), shown in Figure 6C) contained both histidine and glutamine in the Ni^2+^ binding regions where glutamines were either directly interacting (despite weakly) with the Ni^2+^ ion (4ppt) or located in a close proximity to the histidine that attracted the Ni^2+^ (5ib4 and 6ei2). Nonetheless, the roles of these glutamine remain unclear.

## DISCUSSION

We set out to investigate the Ni-NTA binding affinity to various antibody variants in our previous work (Lua et al., 2019) that found different KDs of Pertuzumab and Trastuzumab VH IgE variants interacting with Ni-NTA His-tagged FcεRIα. It is from these data that we serendipitously found that some of the IgE variants were able to bind Ni-NTA.

Other approaches of measuring the IgEs interaction with FcεRIα, such as protein-L mobilized Pertuzumab VH3 IgE to FcεRIα, were not successful (Supplementary Figure S4), restricting the experimental approach to still utilize Ni-NTA binding of His-tagged FcεRIα. It was thus serendipitously found that some variants of the recombinant Pertuzumab and Trastuzumab could bind Ni-NTA. Pertuzumab VH5 IgE bound to FcεRIα with a distinct low KD, i.e. ∼0.066 nM shown in Figure 3 of our previous work (Lua et al., 2019), while the other VH families of Pertuzumab IgE did not show any significant differences from the Pertuzumab VH3 IgE control. This suggests that certain Ni-NTA interactions via the sensors might have contributed into the resulting better KD of the Pertuzumab VH5 IgE variant. Nonetheless, for the low KD of the Trastuzumab VH1, VH3, and VH7 IgE variants (Figure 4 of Lua et al., 2019), the differences with respect to the FcεRIα binding were not confounded by the Ni-NTA sensors because Trastuzumab VH3, VH5, and VH7 IgE variants exhibited similar KD when binding the Ni-NTA. Similarly, the Pertuzumab VH6 IgE binding to FcεRIα (despite high KD, Figure 3 of Lua et al., 2019) was also free from the Ni-NTA interference due to its poor response to the Ni-NTA binding. Although some Ni-NTA interferences on the FcεRIα binding of the Pertuzumab VH5 IgE was found, the consequence did not undermine the conclusion that the VH of IgE can affect FcεRIα binding observed in our previous study.

Further investigation varying the FWRs and/or CDRs of Pertuzumab and Trastuzumab IgE and IgG1 could confer additional Ni-NTA binding functions. Trastuzumab variants (except for VH5-Vκ1 IgE) bound stronger (with lower KD) to the Ni-NTA when compared to the Pertuzumab counterparts as both IgE and IgG1. Through measurements with other heavy metals (Cu^2+^ and Co^2+^) and NTA alone, we ruled out interactions with NTA, narrowing the interactions specifically to the metals. The incidental ability of therapeutic antibodies to bind metals may therefore lead to adverse effects in clinical trials that would need to be avoided during holistic design as discussed in (Ling et al., 2020).

Switching the heavy chain CH between IgE and IgG1 also did not restore the Ni^2+^ binding (abolished in the case of Pertuzumab VH3 IgE), demonstrating a role for the distal CH in Ni binding. The CH exhibited little Ni-NTA binding in most of the Pertuzumab VHs IgG1 variants, a finding in agreement with previous findings (Rosen et al., 2014) despite the presence of the known metal-binding histidine cluster “HEALHNH” (Hale and Beidler, 1994) in IgG1 CH.

While histidine is the canonical binding residue to Ni^2+^, we found that residue numbers alone did not determine the Ni^2+^ binding (unlike the use of 6X His for protein purifications). Among the VH/Vκ families and germlines found in the IMGT (Giudicelli et al., 2005), only a few of VH3, VH4, Vκ1 (Supplementary table S3 and S4), and VH5 framework contained an extra histidine, and they represent only a minority in the overall antibody population (Tiller et al., 2013). Given that IgE is present in much lower quantities than IgGs in human blood (Gould et al., 2003), Ni allergy via immediate IgE encounter due to the extra histidine is very unlikely.

We demonstrated that Ni binding could be modulated with differential pairing of Vκ families while maintaining the histidine content in the sequence. With this, we were able to narrow the binding regions to glutamine stretches (continuous clusters of at least 3 glutamine residues) that were formed spatially with respect to structural arrangement of domains, e.g. at the heavy and light chain interfaces of the Trastuzumab IgE variants or at scaffolds between Vκ1 FWR1 and CDR3 of the Pertuzumab VH5 IgG1 variants. Such an observation lends support to the need for a holistic analysis (Phua et al., 2019) of antibodies, especially when targeting for therapeutic use (Ling et al., 2020) to ensure that therapeutic antibodies do not incidentally cause Ni-induced side effects. These molecular insights of the Ni^2+^ binding in the IgE variants may underlie the pathogenesis of type I nickel hypersensitivity. A number of our IgE variants exhibited better Ni^2+^ binding than their IgG1 counterparts, having implications in AllergOncology. Similarly, this also gives rise to antibody engineering methods to induce Ni^2+^ binding for biotechnological purposes.

Certain associations between Type I and IV hypersensitivity (Gelardi et al., 2017; Kolberg et al., 2020) with a particular clinical case having both underlying Ni allergies (Walsh et al., 2010) warrant further investigation. Since hapten-specific T cells (Saito et al., 2016) in type IV hypersensitivity can involve Ni^2+^ bound MHC-peptide complexes (Moulon et al., 1995; Romagnoli et al., 1991; Sinigaglia et al., 1985), the Ni^2+^ ion could bridge a human leukocyte antigen (HLA-DR) β chain and the hypervariable region of the TCR α chain via interaction with a histidine residue of the HLA-DR (Gamerdinger et al., 2003). However, we did not find structural support for those observations, although a recent report showed that the peptide-bound HLA-B*27:05 complex (PDB: 5ib4) could bind Ni^2+^ ions and the phenomenon was associated with rheumatic disorders (Driller et al., 2019). This peptide had an underlying Ni^2+^ induced conformational reorientation, likely initiating varying T cell responses via an energetically driven mechanism required for the Ni^2+^ binding stabilization. This is similar to the Ni^2+^ binding (via Ni-NTA) to the H67 residue of the Pertuzumab and Trastuzumab VH5-Vκ1 IgE, where the Ni-NTA binding possibly occurred at the H67 in the former, but not in the latter. Furthermore, our recombinant IgE variants maintained Her2 specificity (Lua et al., 2019), and the Ni-NTA binding did not notably compromise the binding to Her2, suggesting that Ni^2+^ binding did not occur at the CDRs, ruling out nickel-switch effects towards the antigen binding as in the engineered dual-specific VHH anti-RNase antibody (Fanning et al., 2014).

Since metal binding can influence antigen presentation of both MHC class I and II (Driller et al., 2019; Wang and Dai, 2013), potentially modulating TCR recognition towards Type IV hypersensitivity, the non-antigenic mechanism of Ni^2+^ binding (also found in our IgE variants via Ni-NTA) may underlie both nickel Type IV and I hypersensitivity pathogenesis, especially in the clinical case reporting both the representations (Walsh et al., 2010). We showed that Pertuzumab VH5-Vκ1 IgE could clearly bind multiple metals via the metal-protein interaction mechanism, supporting that protein-metal interaction (Hemdan et al., 1989) underlies the binding mechanism in antibody as reported in some clinical allergy studies (Bright et al., 1997; Novey et al., 1983; Sastre et al., 2001; Shirakawa et al., 1990), and that these interactions are not as specific as antigen binding (Sela-Culang et al., 2013). The IgE variant might have involved both its glutamine and histidine in binding the Ni^2+^. While glutamines were also found to exhibit long-range effects on protein-metal complex stability it may intervene as a possible binding on-off switch between Ni and of the other metal ions, paving a way for proposing an underlying modulation for the multiple metal allergy that requires further investigation. With such incidental functions, therapeutic antibodies with the unintentional structural stretches of glutamines could confer multi-metal induced allergies (depending on the isotype or immune protein where IgE for type I and HLA for type IV) adverse side effects could arise.

Yet, in all these, given that Ni has been found to an adjuvant to other metals (Bonefeld et al., 2015), there may be a chance for it to be used in vaccines for diseases, e.g. SARS-CoV-2 in (Sekine et al., 2020), that would benefit from a strong T-cell mediated immunity over a humoral one. Further research in tailoring Ni towards such uses would be of interest, but also bearing in mind of inducing Ni hypersensitivity.

In summary, our investigation of recombinant IgEs to His-tagged proteins serendipitously revealed an incidental gain of Ni-binding ability by antibodies without observable compromise to the original antigen. From our analysis, we were able to decipher this gain of function to be contributed by the CDR, FWRs in both VH and Vκ of the antibody with effects from the CH in forming structural stretches of glutamines. This finding may underlie the pathogenesis of type I and type IV nickel hypersensitivities and may be a feature with relevant for both antibody production as well as preventing undesirable adverse effects in therapeutic antibodies.

## ACKNOWLEDGEMENT

This work was supported by A*STAR core funding.

## AUTHOR CONTRIBUTIONS

WHL, WLL, and JJP produced the recombinant antibodies and performed the biolayer interferometry experiments. JY performed the framework germline experiment. CTTS performed the computational simulation and analyses. CTTS, WHL, and SKEG analyzed the results and wrote the manuscript. SKEG designed and supervised all aspect of the study. All authors read and approved the manuscript.

## STAR Methods

### LEAD CONTACT AND MATERIALS AVAILABILITY

Further information and request for resources and reagents should be directed to and will be fulfilled by the Lead Contact, Samuel Ken-En Gan.

### EXPERIMENTAL MODEL AND SUBJECT DETAILS

EXPI293F (human embryonic kidney, GIBCO) cells were cultured in high glucose Dulbecco’s Modified Eagle Medium (DMEM, GIBCO) containing 1X Penicillin-Streptomycin (Nacalai Tesque), 2 mM L-Glutamine (Biological Industries), and 10% heat-inactivated FBS (GIBCO), at 37°C and 5% CO2.

## METHOD DETAILS

### Production and purification of antibodies

Pertuzumab/Trastuzumab IgE VH families variants and Pertuzumab/Trastuzumab IgG1 VH/Vκ families variants were produced as previously described in our previous studies (Lua et al., 2019) and (Ling et al., 2018), respectively. We also produced the Pertuzumab and Trastuzumab VH5/VH3 IgG1 variants that paired with different Vκ families (Vκ1 to Vκ6) for investigation of Ni-NTA binding affinities in the same manner. We used protein L to purify all the IgE variants while using protein G column for IgG1 variant purification.

### Bio-layer interferometry studies

Measurements of the association and dissociation rate constants of the antibodies to the Ni-NTA were carried out by directly binding the antibodies at concentrations with two-fold serial dilutions from 200 nM down to 3.125 nM to NTA biosensors (Fortebio) recharged with 10mM NiCl_2_ for 1 min. All measurements were performed using the 1× kinetic buffer (Fortebio) on the Octet Red96 system (Fortebio).

For cross metal reactivity assay, Ni-NTA biosensors were stripped with low pH 1.7 10mM Glycine and recharged with 10mM of Cobalt (II) Chloride (CoCl_2_), Nickel (II) Chloride (NiCl_2_), or Copper (II) Chloride (CuCl_2_), before binding with 100nM of Pertuzumab VH5 IgE variants.

### Structural modeling of full length Trastuzumab and Pertuzumab VH3, VH5, and VH7 IgE and VH5 IgG1 variants

Models of the full length Pertuzumab and Trastuzumab VH3, VH5, and VH7 IgE paired with their respective Vκ1 were constructed previously (Lua et al., 2019). To obtain the Pertuzumab VH5-Vκ1 IgG1 variant, we replaced the IgE CH of the Pertuzumab VH5-Vκ1 IgE structure with the IgG1 CH. The resulting Pertuzumab VH5-Vκ1 IgG1 model was minimized and equilibrated (1 nanosecond or ns, explicit solvent with sequential constant volume and pressure, NVT and NPT ensembles, respectively) using GROMACS 2019.3 (Abraham et al., 2019).

Similarly, by replacing the light chain V-region (Vκ1) with Vκ3 and Vκ6 of the full length Pertuzumab and Trastuzumab VH5 IgG1 (excluding the others for compatible comparison with experimental results), we obtained structural models of those VH5 IgG1 variants for analyzing various effects of different Vκ.

### Dynamics simulation of the Trastuzumab and Pertuzumab IgE^Fab^ and IgG1^Fab^ variants

We extracted the Fab regions of all the full length models of Pertuzumab and Trastuzumab IgE and IgG1 variants to further investigate the dynamics of the VH-Vκ pairing.

The Fab structure was first capped with ACE/NME at the N- and C-terminus, respectively, to maintain the neutral charges and avoid artificial interference at the two termini of each single Fab domain. The system was then relaxed by subjecting it to short minimization (5000 steps) and heated in gradual thermal baths from 0-100K (with constant volume) and then from 100-300K (with constant pressure). The system was then equilibrated (1 ns) and followed by 100 ns production applying explicit solvent model in triplicates (3×100 ns). The molecular dynamics (MD) simulations were carried out with random velocities and constrained by the Langevin temperature equilibration scheme to thermalize the systems at 300K at time steps of 2 fs, using AMBER14 (Case et al., 2015).

### Blind docking of the monomer parameterized nickel (Ni^2+^)-bound NTA onto the Pertuzumab and Trastuzumab VH3, VH5, and VH7 IgE^Fab^ domains

The Ni-bound NTA (nitrilotriacetic acid) was first constructed using topologies of NTA (from PDB: 1gvc) and of Ni^2+^ ion (from PDB: 4ppt). AM1-BCC charge method (implemented in antechamber) was applied, using Chimera (Pettersen et al., 2004), to compute partial charges of the constructed Ni-bound NTA, thereafter name Ni-NTA.

Blind dockings using Autodock Vina (Trott and Olson, 2010) were performed on the resulting Ni-NTA (as ligand) and the Pertuzumab and Trastuzumab VH3, VH5, or VH7 IgE^Fab^ domains (since the Fc domain is inferred to be weakly involved in the nickel binding in our experiments) to explore searching space for possible Ni-NTA binding. Bonds in the Ni-NTA were kept rigid to minimize the confounded binding (if any) of the NTA to the respective Fab domain (as receptor) while also constraining the flexibility of the free Ni^2+^ from binding to the deeply buried cavities. A 1-Å grid box was placed at the molecule center and extended to cover the whole Fab domain. Pre-defined parameters of Ni^2+^ ion in the AutodockTools packages were used. The dockings were performed for 10 replicates that represent 10 different IgE^Fab^ conformations of each variants, resulting 10×1000 conformers for each, and with exhaustive searching (exhaustiveness level set at 128). A distance cutoff of 4 Å between the Ni^2+^ ion of the Ni-NTA and the interacting residue was used to determine the binding.

### Cryptic pocket detection

Cryptic binding site detection was performed on Fab domains of Pertuzumab and Trastuzumab VH3-Vκ1, VH5-Vκ1, or VH7-Vκ1 IgE, using an online server CryptoSite (Cimermancic et al., 2016). The resulting predicted cryptic residues were then matched into identified pockets that were resulted from a concurrent pocket detection, using *mdpocket* (Schmidtke et al., 2010) on the MD simulation trajectories. Only internal (hidden) pockets or open channels (with isovalue ≥ 5) that were detected for more than 80% of the trajectories were chosen and defined as dense (isovalue ≥ 8) or less dense (5 ≤ isovalue < 8) cavities. If the chosen internal pocket involves at least one of the predicted cryptic residues by CryptoSite, it is considered as a cryptic pocket.

### Structural analysis of protein-metal complexes from Protein Data Bank (PDB)

We searched protein-metal complexes of Ni^2+^, Cu^2+^, and Co^2+^ available from PDB (accessed in June 2020) by using the advance ligand search function in the PDB with NI, CU, or CO as keywords in the “Identifier” search. Only those protein complexes that contain the metal as standalone ligand were retrieved. We further narrowed the search to look for protein complexes that were involved in the immune system, by extending the keywords to “immune”, “antibody”, “MHC”, and “TCR”.

### QUANTIFICATION AND STATISTICAL ANALYSIS

All dissociation equilibrium constant measurements were conducted with at least duplicate measurements at various concentrations from 200 nM to 3.125 nM of the antibodies. Values of KD (M), ka (1/Ms), and kd (1/s) were measured and calculated using the Octet RED96 system and analysis software. “Poor response” is determined for the cases where the responses are less than 0.1 nm for at least three concentrations. Graphs of the representative sets were plotted using Microsoft Excel 2010.

### DATA AND CODE AVAILABILITY

All data are available in the manuscript or the Supplemental Information while raw datasets are available from the corresponding author on reasonable request.

## SUPPLEMENTARY FIGURES

**Figure S1.**
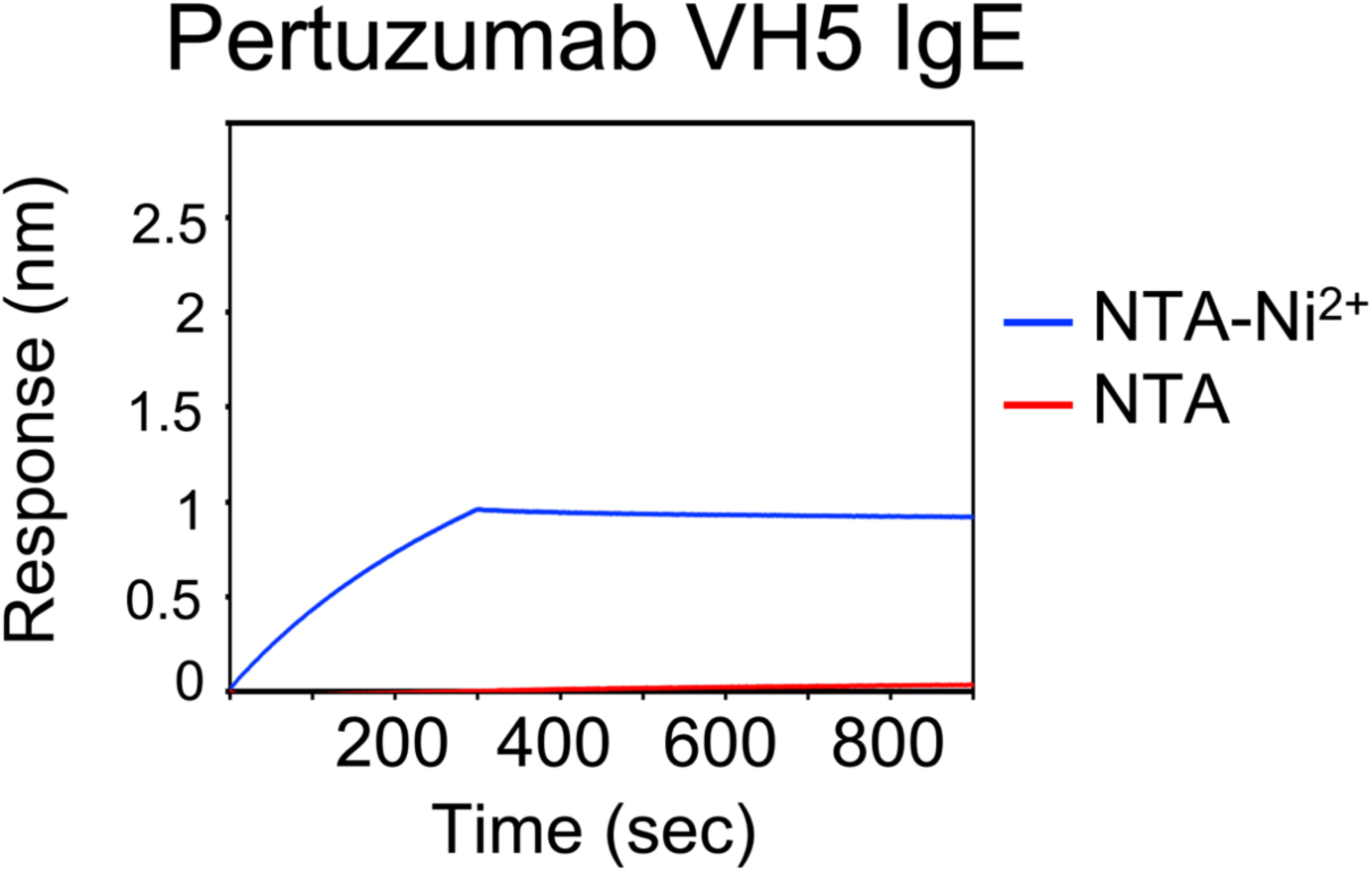
Binding response of the Pertuzumab VH5 IgE variant to the Ni-NTA sensor that was recharged with NiCl_2_ (NTA-Ni^2+^) versus NTA with only water (NTA) to rule out interactions with the NTA sensor. The result shown here is a part of those presented in Figure 6A.

**Figure S2:**
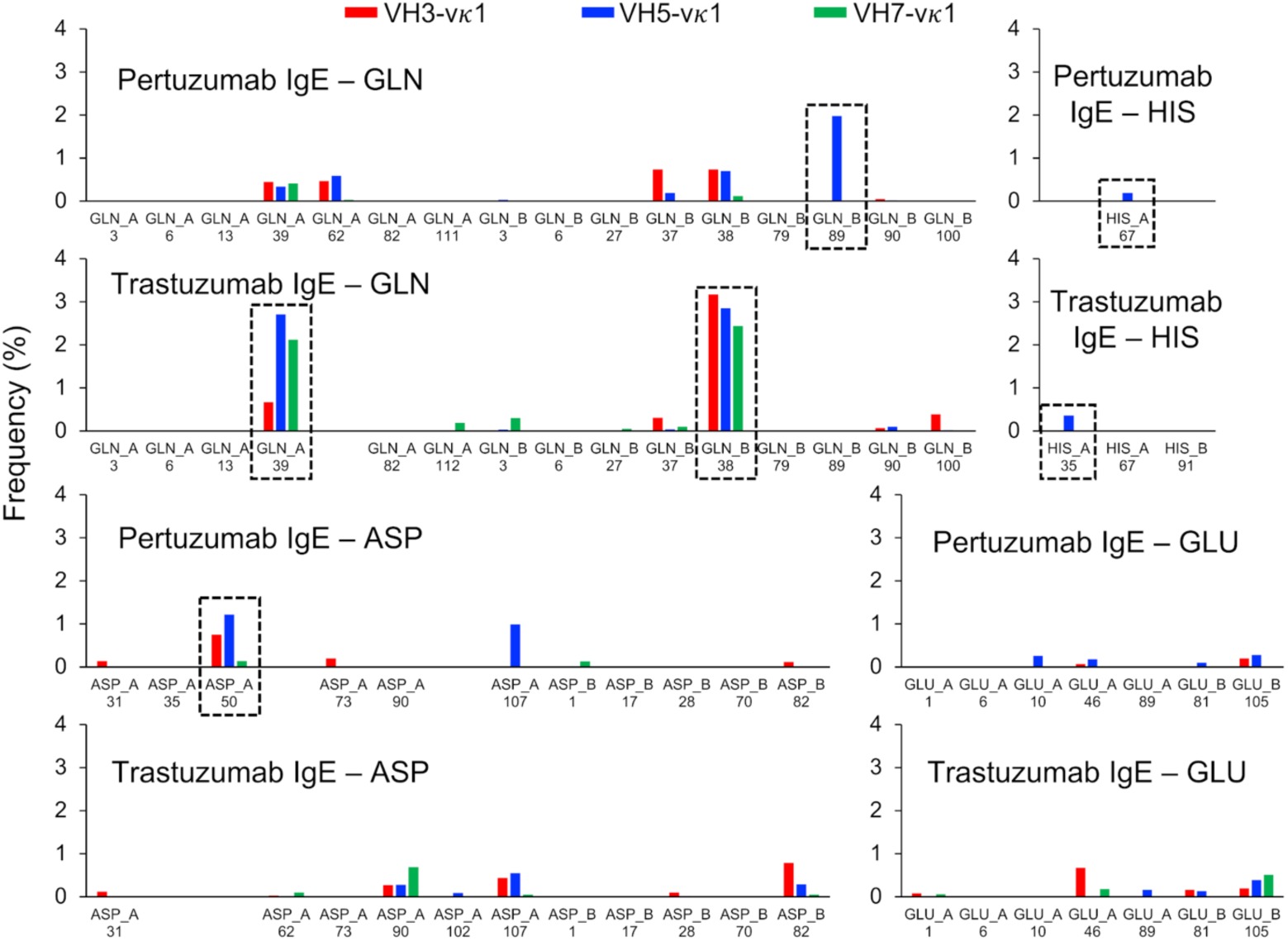
Distribution of the Ni-NTA conformers clustered at the residues of interest such as glutamine (GLN), histidine (HIS), aspartate (ASP), and glutamate (GLU) on the Pertuzumab and Trastuzumab IgE V-regions of VH3, VH5, and VH7 variants. No measurements were readable at cysteine residues. Predominant clusters of the Ni-NTA conformers are highlighted in dashed boxes. Residues in the heavy and light chains are labeled as chain A and B, respectively.

**Figure S3:**
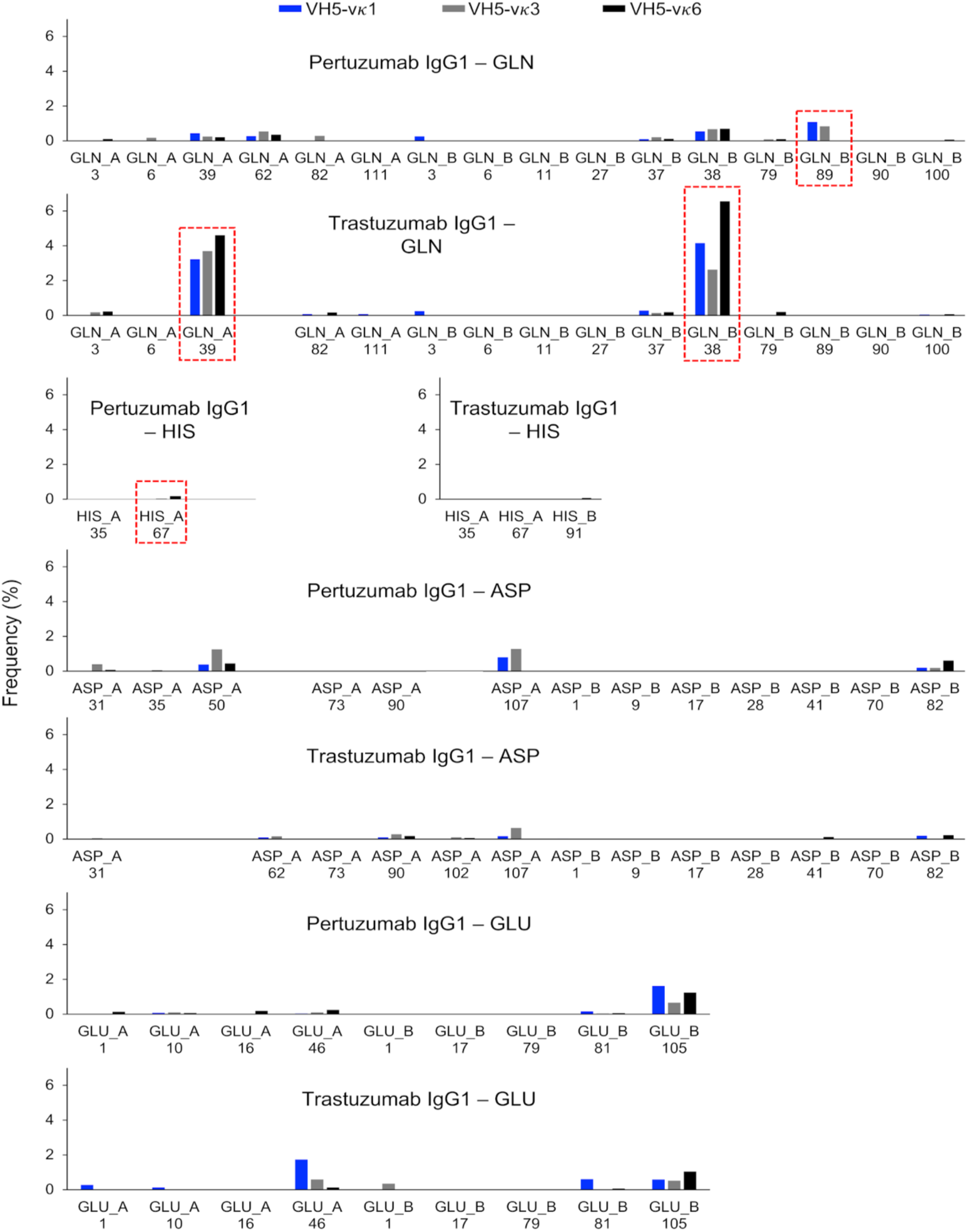
Distribution of the Ni-NTA conformers clustered at the residues glutamine (GLN), histidine (HIS), aspartate (ASP), and glutamate (GLU) on the Pertuzumab and Trastuzumab IgE V-regions of VH5 paired with Vκ1, Vκ3, and Vκ6. The clusters of interest of the Ni-NTA conformers are highlighted in dashed boxes. Residues in the heavy and light chains are labeled as chain A and B, respectively.

**Figure S4.**
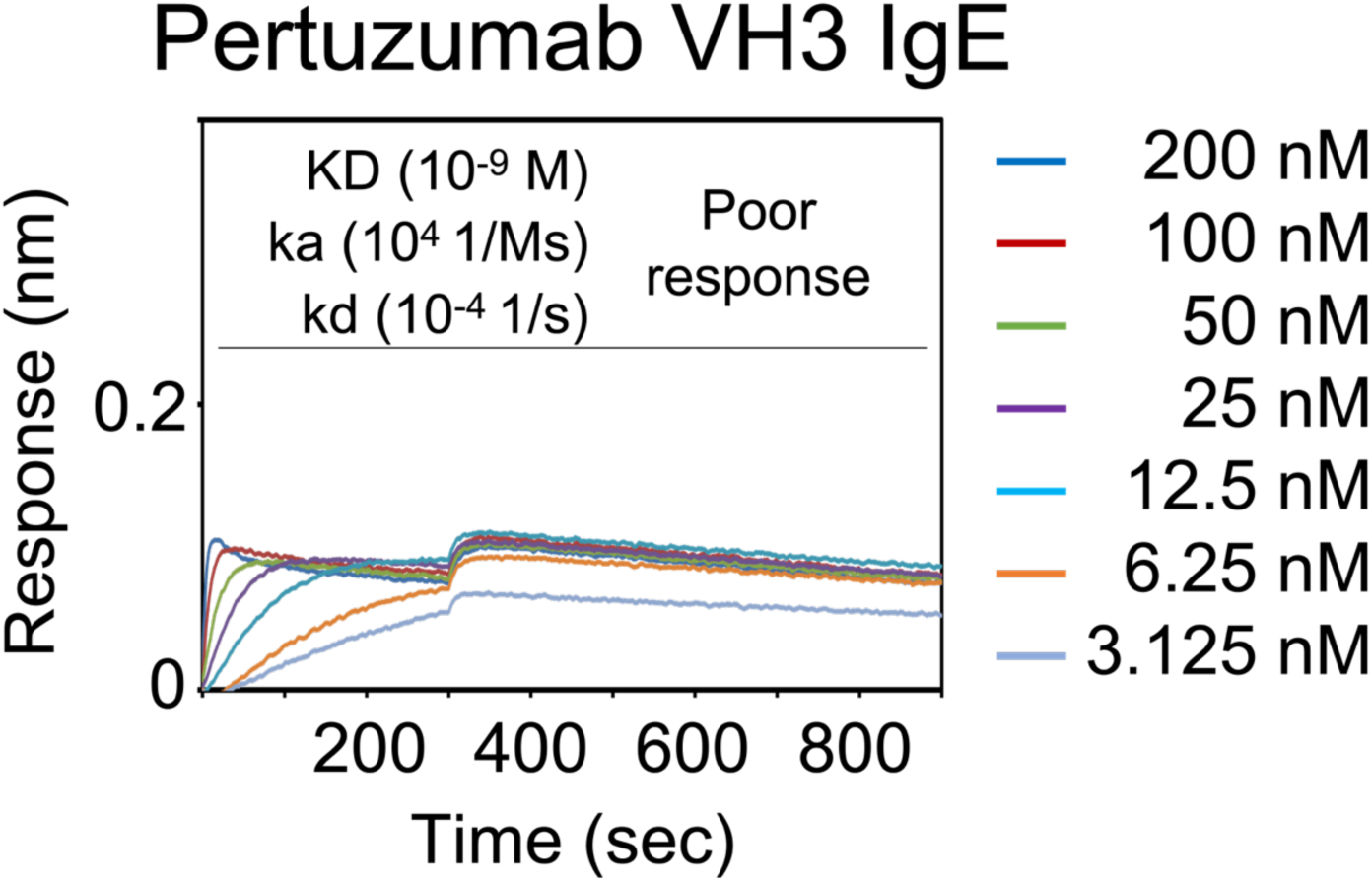
Dissociation equilibrium constants (KD) of the FcεRIα to the immobilized Pertuzumab VH3 IgE (using Protein L biosensor). Biolayer interferometry measurements of the FcεRIα at various concentrations from 200nM to 3.125nM. Values of KD (M), ka (1/Ms), and kd (1/s) were measured and calculated using the Octet RED96 system. The X-axis depicts the time (in seconds) and the Y-axis depicts the binding response (nm).

**Supplementary Table S1:** Residue content of Histidine, Glutamine, Glutamate, Aspartate, and Cysteine present in the antibody variants of the study. Where the variants share the same domain, only one value is presented for that domain count. For examples, all Pertuzumab (or Trastuzumab) VHs (or Vκs) variants share the same heavy chain (or light chain) CDRs, respectively. All IgE (or IgG1) variants share the same heavy chain constant region (CHs). Only Cκ is used as light chain constant region in this study.

|  |  | Pertuzumab / Trastuzumab |  |  |  |  |  |  |  |  |  |  |  |  |  |
| --- | --- | --- | --- | --- | --- | --- | --- | --- | --- | --- | --- | --- | --- | --- | --- |
|  |  | Heavy Chain |  |  |  |  |  |  | Light Chain |  |  |  |  |  |  |
|  |  | VH1 | VH2 | VH3 | VH4 | VH5 | VH6 | VH7 |  | Vk1 | Vk2 | Vk3 | Vk4 | Vk5 | Vk6 |
| **HIS (H)** | H-FWRs | 0 / 0 | 0 / 0 | 0 / 0 | 0 / 0 | 2 / 2 | 0 / 0 | 0 / 0 | L-FWRs | 0 / 0 | 0 / 0 | 0 / 0 | 0 / 0 | 0 / 0 | 0 / 0 |
|  | H-CDRs |  |  |  | 0 / 2 |  |  |  | L-CDRs |  |  | 0 / 2 |  |  |  |
|  | CHs |  |  | 22 (IgE) | – | 18 (IgG1) |  |  | Ck |  |  | 4 |  |  |  |
| **GLN (Q)** | H-FWRs | 16 / 14 | 12 / 10 | 12 / 10 | 18 / 16 | 12 / 10 | 20 / 18 | 12 / 10 | L-FWRs | 12 / 12 | 10 / 10 | 10 / 10 | 12 / 12 | 10 / 10 | 12 / 12 |
|  | H-CDRs |  |  |  | 2 / 0 |  |  |  | L-CDRs |  |  | 6 / 6 |  |  |  |
|  | CHs |  |  | 38 (IgE) | – | 22 (IgG1) |  |  | Ck |  |  | 12 |  |  |  |
| **GLU (E)** | H-FWRs | 6 / 6 | 4 / 4 | 8 / 8 | 2 / 2 | 8 / 8 | 4 / 4 | 8 / 8 | L-FWRs | 4 / 4 | 8 / 8 | 10 / 10 | 6 / 6 | 10 / 10 | 10 / 10 |
|  | H-CDRs |  |  |  | 0 / 0 |  |  |  | L-CDRs |  |  | 0 / 0 |  |  |  |
|  | CHs |  |  | 38 (IgE) | – | 34 (IgG1) |  |  | Ck |  |  | 14 |  |  |  |
| **ASP (D)** | H-FWRs | 8 / 6 | 8 / 6 | 6 / 4 | 6 / 4 | 6 / 4 | 6 / 4 | 6 / 4 | L-FWRs | 8 / 8 | 8 / 8 | 4 / 4 | 10 / 10 | 6 / 6 | 8 / 8 |
|  | H-CDRs |  |  |  | 8 / 8 |  |  |  | L-CDRs |  |  | 2 / 2 |  |  |  |
|  | CHs |  |  | 36 (IgE) | – | 26 (IgG1) |  |  | Ck |  |  | 10 |  |  |  |
| **CYS (C)** | H-FWRs | 4 / 4 | 4 / 4 | 4 / 4 | 4 / 4 | 4 / 4 | 4 / 4 | 4 / 4 | L-FWRs | 4 / 4 | 4 / 4 | 4 / 4 | 4 / 4 | 4 / 4 | 4 / 4 |
|  | H-CDRs |  |  |  | 0 / 0 |  |  |  | L-CDRs |  |  | 0 / 0 |  |  |  |
|  | CHs |  |  | 26 (IgE) | – | 18 (IgG1) |  |  | Ck |  |  | 6 |  |  |  |

**Supplementary Table S2:**
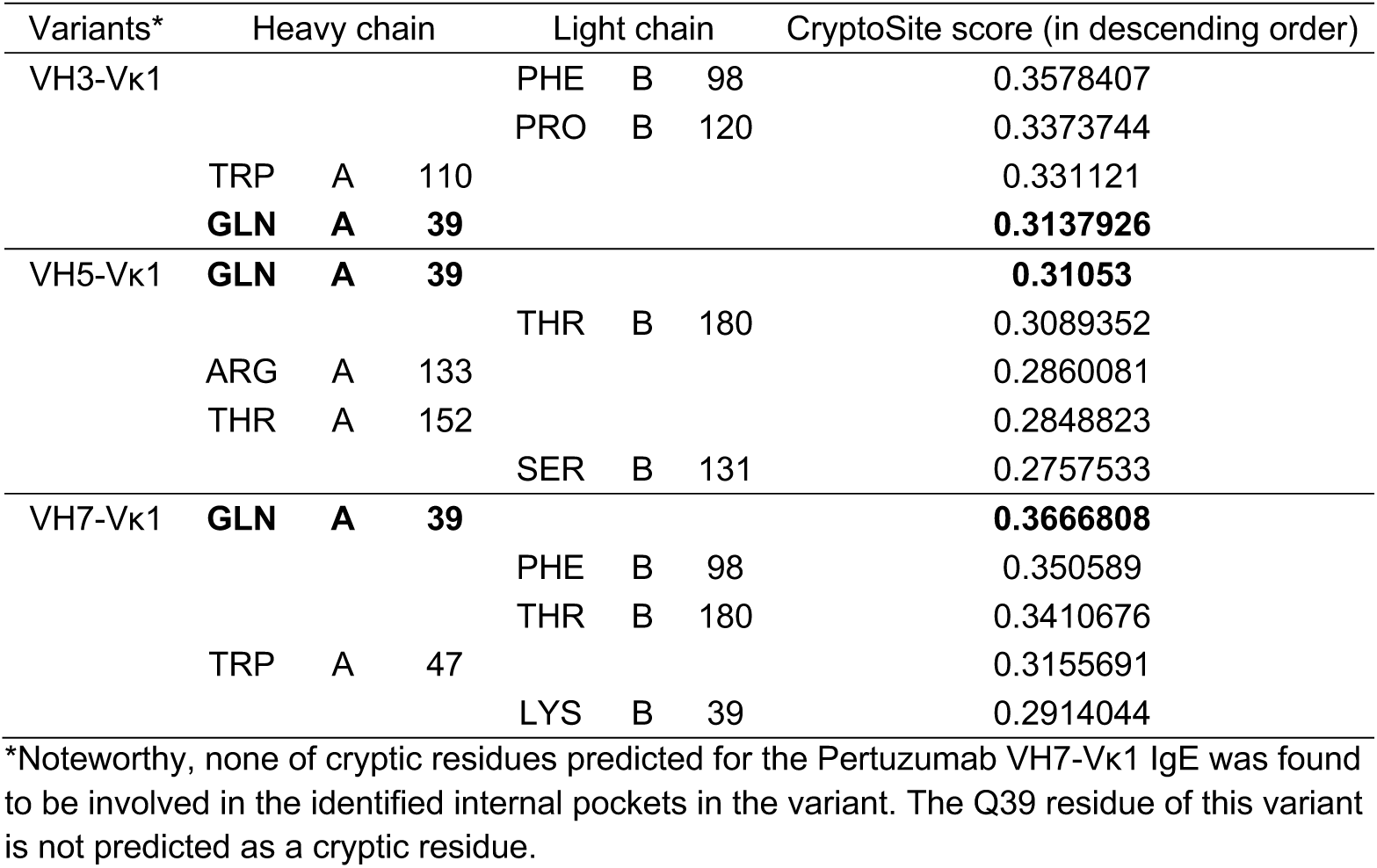
Top predicted cryptic residues of the Trastuzumab VH3-Vκ1, VH5-Vκ1, and VH7-Vκ1 IgE using CryptoSite (Cimermancic et al., 2016). The difference cutoff of scores between two consecutive residue results is 0.05 to determine the selected residues from the first highest score. The Q39 residues that were matched in the identified internal pockets are in bold.

**Supplementary Table S3:** Antibody framework repertoire analysis. Analysis of different light chain framework family germline for presences of histidine and estimated germline occurrence (Tiller et al., 2013)

| Families / Germline | Extra histidine | Frequency (%) |
| --- | --- | --- |
| Vk1-5 | - | 11 |
| Vk1-6 | - | >1 |
| Vk1-8 | - | 2 |
| Vk1D-8 | - | >1 |
| Vk1-9 | - | >1 |
| Vk1-12 | - | >1 |
| Vk1-13 | - | >1 |
| Vk1D-13 | - | >1 |
| Vk1-16 | - | >1 |
| Vk1-17 | + | >1 |
| Vk1-27 | - | 2 |
| Vk1-33 | - | 2 |
| Vk1D-33 | - | 2 |
| Vk1-39 | - | 7 |
| Vk1D-39 | - | 7 |
| Vk1D-43 | - | >1 |
| Vk2 – ALL | - | 11 |
| Vk3 – ALL | - | 36 |
| Vk4 – ALL | - | 8.5 |
| Vk5 – ALL | - | >1 |
| Vk6 – ALL | - | >1 |

**Supplementary Table S4:**
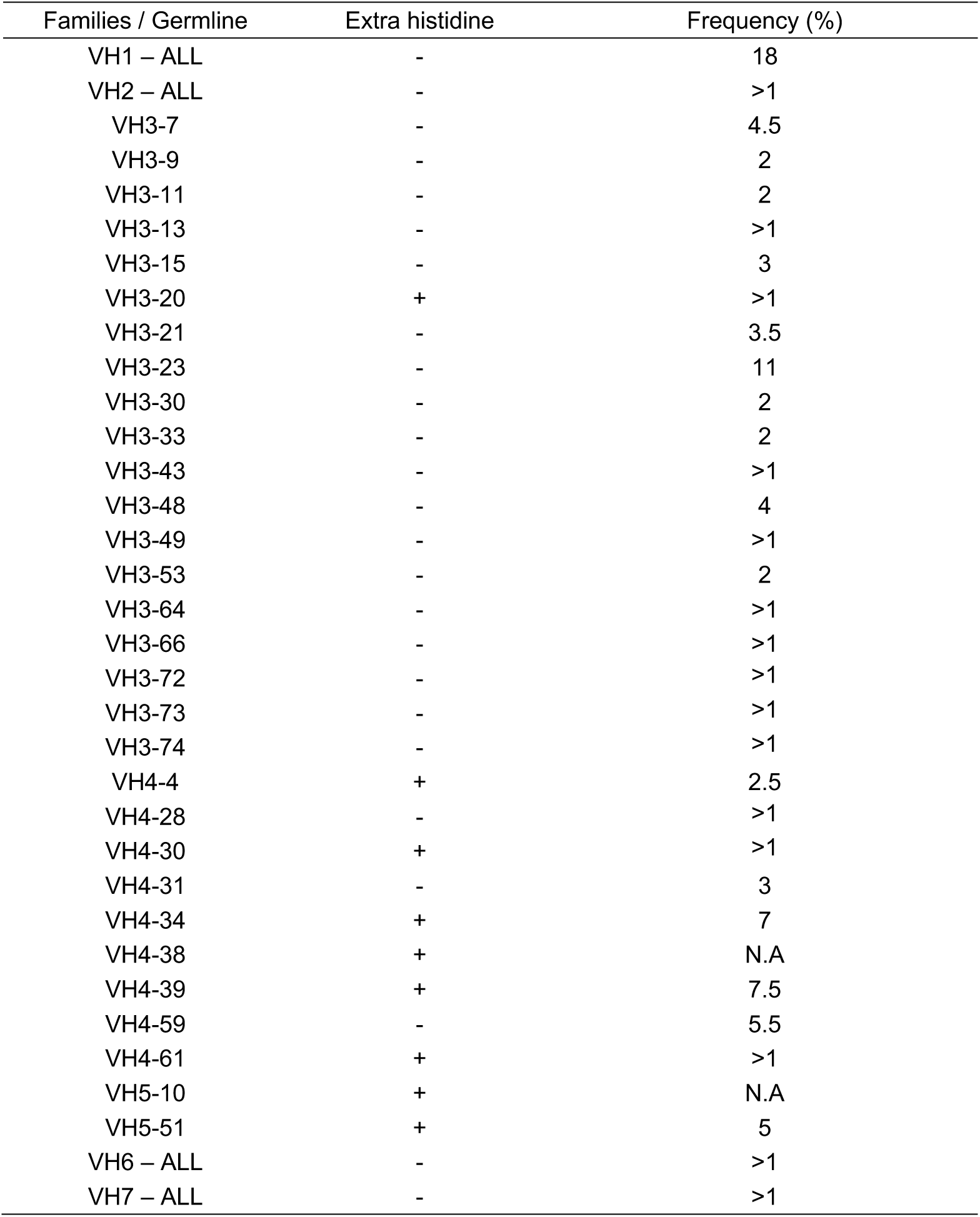
Antibody framework repertoire analysis. Analysis of different heavy chain framework family germline for presences of histidine and estimated germline occurrence (Tiller et al., 2013)

